# A reproducible buccal transcriptomic response to music exposure in autism spectrum disorder: insights into the oral-brain axis

**DOI:** 10.64898/2026.09.07.749867

**Authors:** Alberto Gómez-Carballa, Nour El Zahraa Mallah, Laura Navarro, Federico Martinón-Torres, Antonio Salas

**Author notes:** Corresponding authors: Antonio Salas.

## Abstract

Music-based interventions (MIs) have shown promise in autism spectrum disorder (ASD), yet the molecular mechanisms underlying their effects remain poorly understood. The transcriptomic response to music exposure in individuals with ASD and healthy controls (HCs) was analyzed using buccal swab samples and a targeted NanoString gene expression panel. The discovery cohort included baseline and post-intervention samples, whereas the validation cohort added a 25-minute intermediate sample to assess transcriptional dynamics. Thirty-three candidate genes were identified in the discovery cohort, of which 20 (61%) remained significantly differentially expressed in the independent validation cohort, including 13 that withstood multiple-testing correction. Replicated genes included several key regulators of innate immunity and inflammatory signaling, such as *CXCL8*, *MYD88*, *RELA*, *PTGS2*, *PTPRC*, *SELL*, *LYN*, and *LTBR*. Functional enrichment analyses revealed robust innate immune and inflammatory pathways, involving TLR/IL-1-MYD88 signaling, cytokine activity, leukocyte activation, and antimicrobial responses. Co-expression network analysis identified coordinated remodeling of immune transcriptional programs in ASD. Consensus feature selection identified a two-gene signature (*FURIN* and *JUNB*) that accurately discriminated pre- and post-musical stimulation samples in ASD but not in HCs. The same signature detected transcriptomic responses after only 25 minutes of music exposure by discriminating baseline and intermediate samples, with intermediate expression levels between baseline and post-intervention values, suggesting a progressive time-dependent response. These findings support a reproducible transcriptional response to music in ASD across independent cohorts implicating neuroimmune pathways. The resulting transcriptional signature supports the development of minimally invasive biomarkers for monitoring MIs in neurodevelopmental disorders.

## Introduction

Autism Spectrum Disorder (ASD) is a neurodevelopmental condition that affects how individuals perceive the world, often involving altered sensory processing, social communication difficulties, and restricted and repetitive behaviors, alongside strengths in diverse domains (Randell et al., 2019; Tavassoli et al., 2016). The impact of ASD can be significant, not just for children but also for their families, influencing daily routines, finances, physical health, and mental well-being (Hossain et al., 2020). Given the multifactorial genetic, developmental, and environmental risk factors associated with ASD, epigenetic mechanisms may contribute to its development (Hall & Kelley, 2014). Current ASD treatments primarily target behavioral symptoms, but many interventions are limited, lack robust empirical validation, and require prolonged treatment. Moreover, their safety and effectiveness remain insufficiently supported by evidence (Randell et al., 2019; S. R. Sharma et al., 2018; Voss et al., 2019; Warren et al., 2011; Wood et al., 2020). Hence, effective therapeutic approaches to improve behavioral outcomes are needed.

Music-based interventions (MIs), including singing, instrumental performance, music listening, and music therapy, are increasingly recognized as promising approaches for individuals with ASD (Chenausky & Schlaug, 2018; Cook et al., 2019). MIs can promote verbal and non-verbal communication, emotional expression, social interaction, and quality of life (Bieleninik et al., 2017; Brown, 2017; Navarro et al., 2025; Quintin et al., 2011; Thompson, 2018). Their accessibility and ability to engage cognitive, emotional, social, and biological processes further support their therapeutic potential.

Musićs benefits extend beyond its fundamental neurological impacts; it plays a crucial role in neuro-rehabilitation practices (Navarro et al., 2025; Pantev & Herholz, 2011). Neuroscience has revealed several connections between music and ASD, particularly highlighting the distinct cognitive and sensory processing characteristics observed in individuals with ASD (Kim et al., 2009; Sharda et al., 2018; Wagener et al., 2021). For example, fMRI demonstrated greater activation of auditory and emotion-processing brain regions in individuals with ASD than in neurotypical controls during music listening, suggesting enhanced neural responsiveness to musical stimuli (Sharda et al., 2018).

Recent evidence from the literature support the idea of that musical stimuli can yield measurable benefits for patients with ASD and other neurodegenerative disorders (Gómez-Carballa et al., 2023; He et al., 2024; Li et al., 2024; Navarro et al., 2023; Navarro et al., 2025; Redondo Pedregal & Heaton, 2021; Särkämo et al., 2008; Sisti et al., 2024; Van de Winckel et al., 2004). Importantly, sensory experiences such as music may also induce molecular changes by modulating gene expression and molecular pathways in biofluids, including saliva (Gómez-Carballa et al., 2025; Gomez-Carballa et al., 2023; Nair et al., 2019; Nair et al., 2021; Navarro et al., 2023), including saliva.

Saliva has emerged as a promising non-invasive biofluid for biomarker discovery owing to its ease of collection and potential to reflect both local and systemic processes (Pfaffe et al., 2011; Roi et al., 2019; V. Sharma et al., 2022). Its autonomic regulation establishes an oral-brain axis linking neural activity with salivary secretion and immune responses (Murai et al., 1998; Sansores-Espana et al., 2021). Accordingly, salivary molecules are increasingly investigated as biomarkers of brain health, stress, and neurological disorders (Bauduin et al., 2021; Schepici et al., 2020). In ASD, salivary profiling has identified biomarkers related to neurodevelopment, synaptic plasticity, and immune regulation, supporting saliva as a valuable matrix for studying disease mechanisms and therapeutic responses (Hicks et al., 2016; Hicks et al., 2018; Janšáková et al., 2021).

The first multi-omics study of musical stimulation in ASD, combining saliva-based transcriptome and microbiome analyses, identified coordinated changes in genes and pathways related to neurodevelopment, immune regulation, energy metabolism, and inflammation (Cavenaghi et al., 2025). These findings suggest that music may influence host molecular pathways and host-microbiome interactions. However, the study was a proof-of-concept investigation with a modest sample size, and whole-transcriptome sequencing of saliva was limited by abundant microbial RNA and the low and variable quantity of host RNA (Cavenaghi et al., 2025). In addition, the use of saliva for whole-transcriptome RNA sequencing introduced technical limitations related to the complexity of the salivary transcriptome. Saliva contains substantial amounts of microbial RNA, which can represent a considerable proportion of the total RNA pool and consequently reduce the fraction of sequencing reads attributable to human transcripts, thereby limiting the depth and breadth of host transcript detection (Gosch et al., 2024).

To address these limitations, the present study provides a more focused and scalable assessment of host gene-expression responses to musical stimulation in ASD. We included a larger sample of TEA cases and healthy controls (HCs) and evaluated transcriptomic changes before and after brief musical stimulation. A targeted, hybridization-based, PCR-free nCounter assay (NanoString), combined with a dedicated saliva collection device, was used to quantify predefined transcripts relevant to neurodevelopment, immune regulation, inflammation, and ASD-related pathways while minimizing the impact of microbial RNA and variable RNA quality. Building on previous proof-of-concept findings, this study represents, to the best of our knowledge, the first investigation of music-associated salivary transcriptomic changes in autistic individuals and HCs, offering a novel approach to characterize host molecular responses to musical stimulation in ASD.

## Methods

### Experimental design and sample collection

Under the framework of the Sensogenomics project (https://sensogenomics.com), which investigates the biomolecular and physiological effects of music in different human populations (Gómez-Carballa et al., 2025; Navarro et al., 2021; Salas et al., 2025), two experimental concert studies were conducted at the Auditorio de Galicia (Santiago de Compostela, Galicia, Spain). The first cohort (Sensogenoma22) was recruited in 2022 and served as the discovery cohort, whereas the second cohort (Sensogenoma25), recruited in 2025, served as an independent validation cohort. Both concerts were performed by professional musicians from the Real Filharmonía de Galicia, with the 2025 edition additionally featuring the Municipal Band of Santiago de Compostela. All participants remained seated throughout the performances to eliminate the influence of physical activity. The experimental setting was designed to be comfortable, quiet, and non-invasive, thereby minimizing stress and other external factors that could influence physiological responses. The Sensogenoma22 concert consisted of a 50-minute orchestral programme conducted by Baldur Brönnimann, designed to evoke a broad range of emotional responses through variations in timbre, tempo, musical style, and tonality. The repertoire included The Unanswered Question (C. Ives), The Merry Wives of Windsor (O. Nicolai), Slavonic Dances Nos. 2 and 3, Op. 46 (A. Dvořák), Oblivion (A. Piazzolla), Hungarian Dance No. 5 (J. Brahms), The Barber of Seville: Overture (G. Rossini), and Danzón No. 2 (A. Márquez). The Sensogenoma25 concert was deliberately structured to evoke two contrasting emotional states of similar duration (25 minutes). The first half was designed to evoke sadness and included excerpts from Swan Lake (P. I. Tchaikovsky), the third movement of Symphony No. 3 (J. Brahms), Braveheart: For the Love of a Princess (J. Horner), the Intermezzo from Manon Lescaut (G. Puccini), and Nimrod from the Enigma Variations (E. Elgar). Following the intermission, the second half focused on positive emotional responses through performances of selections from The Comedians Suite (D. Kabalevsky), Slavonic Dance No. 3 (A. Dvořák), movements from Romeo and Juliet Suite (S. Prokofiev), and Gayaneh Suite (A. Khachaturian), concluding with Toreadors’ March from Carmen (G. Bizet) as an encore.

All participants were instructed to refrain from eating, drinking, smoking, chewing gum, or engaging in physical activity for at least 30 minutes before saliva collection to ensure the reliability and comparability of transcriptomic analyses.

Saliva samples were collected simultaneously from all participants under the supervision of trained healthcare professionals using Oragene RNA saliva collection devices (ORE-100, DNA Genotek). Participants were instructed to rub the sponge tip of the collection device along both sides of the gums while avoiding contact with the teeth until sufficient saliva had been absorbed. The sponge was then transferred into a tube containing 1 mL of RNA stabilization solution, and samples were stored at room temperature until processing.

The saliva sampling schedule differed between cohorts. In the Sensogenoma22 discovery cohort, samples were collected immediately before the concert (baseline time point, bTP) and immediately after the performance (final time point, fTP). In the Sensogenoma25 validation cohort, an additional sample was collected during the concert intermission (intermediate time point, iTP), resulting in three sampling time points (bTP, iTP, and fTP). **Figure 1** summarizes the overall experimental design.

**Figure 1.**
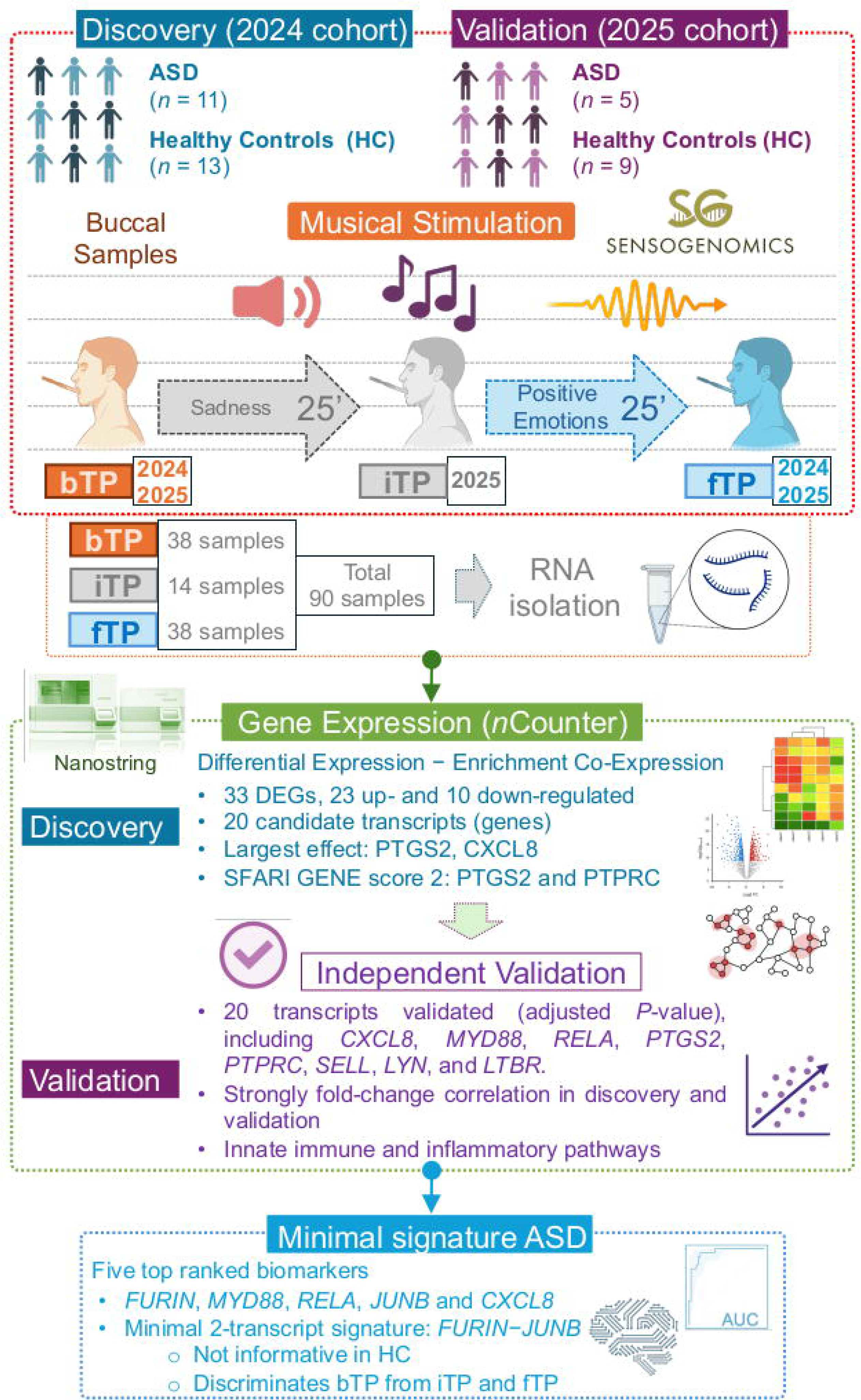
Scheme of the experimental design. The figure was built using Biorender resources (https://www.biorender.com/).

### RNA processing and *n*Counter assay

RNA was isolated from 500 μL of saliva using the RNeasy Micro Kit (Qiagen) following manufacturer-default instructions. Given the sensitivity of transcriptomic analyses to technical variation, extensive efforts were made to minimize potential batch effects. Samples from the two cohorts were processed and analyzed independently, with each cohort analyzed at a different time. Within each cohort, ASD and HC samples were balanced across NanoString cartridges to minimize potential cartridge (batch) effects. Whenever possible, longitudinal samples from the same individual were processed on the same cartridge, although this was not always feasible because of cartridge capacity constraints. All cartridges were processed using identical reagents, protocols, and laboratory conditions. This strategy was adopted to reduce technical variability and maximize confidence that the observed transcriptional differences reflected true biological responses rather than experimental artifacts.

The protocol provided by the extraction kit was slightly modified according to the recommendations from the Oragene tubes supplier. An RNA concentration step and an additional DNase treatment were undertaken using an RNA clean & concentrator kit (Zymo Research). RNA amount and integrity were assessed using TapeStation 4200 (Agilent), calculating DV200 values to confirm that >50% of the RNA fragments were above 200nt, to estimate the optimal sample input.

For gene expression analysis, the *nCounter MAX* (NanoString Technologies) was used in combination with the *n*Counter Host Response Panel, which covers 785 genes. The discovery and validation cohorts were processed and analyzed independently and at different time points. Within each cohort, samples from cases and controls were deliberately mixed and interspersed in different cartridges and processed simultaneously during NanoString gene expression profiling, ensuring that biological groups were not confounded by technical processing effects. Standard protocols were followed, including 12× RNA hybridization with 5 µl of RNA as input and an 18-hour hybridization time for all samples. After excluding genes with expression levels below background (maximum expression value < background threshold), 520 and 501 genes, respectively, were detected in ASD patients and HCs from the 2022 cohort, out of the 785 genes included in the NanoString panel. In the 2025 cohort, 338 genes were detected in ASD patients and 371 in HCs.

### Statistical analysis

A quality control (QC) assessment of the raw expression data was carried out, ensuring to follow the manufacture’s guidelines to avoid any technical issues. Samples that did not meet the QC standards or had a low gene count were excluded. Genes were excluded that had counts falling below the background level, which is defined as the mean + 2 standard deviations (SD) of the negative control spikes in the code set.

For data normalization, an iterative approach that combines the *DEGSeq2* (Love et al., 2014) and *RUVSeq* (Risso et al., 2014) packages was used, as detailed in Bhattacharya et al. (Bhattacharya et al., 2021). To find control reference genes for normalization, invariable genes (with a *P*-value > 0.1, BaseMean > 100, and |log_2_FC| < 0.2) were selected after performing a naïve differential expression analysis between TP1 and TP2 for both ASD patients and HCs separately. Genes expressed below the background were removed. A paired-sampling design to analyze the differences in the transcriptome before (bTP) and after (fTP) the musical stimuli was employed. The discovery and validation cohorts were processed using the same bioinformatic pipeline to ensure a consistent analytical framework across both datasets

To assess the relevance of the identified differentially expressed genes (DEGs) in ASD, all DEGs were cross-referenced against the Simons Foundation Autism Research Initiative (SFARI) Gene database (https://gene.sfari.org/; downloaded 22/06/2026), a curated repository of genes implicated in ASD (Abrahams et al., 2013).

The Weighted Gene Co-expression Network Analysis (*WGCNA*) R package (Langfelder & Horvath, 2008) was used to explore clusters of co-expressed genes that might be correlated with musical stimuli in both ASD patients and HCs, analyzed separately. The analysis began with normalized and corrected gene expression data, which were adjusted for variability among patients, to build a signed weighted correlation network. Following the guidelines from the package developers and considering the number of samples in each group, a soft-thresholding power of 18 was selected. The Topological Overlap Matrix (TOM) and the corresponding dissimilarity values (1-TOM) were then computed. A minimum module size of 30 and a dendrogram cut height threshold of 0.2 were established for merging modules. Initially, the identified modules of co-expressed genes were labeled by colors, but they were later renamed based on the genes with the highest connectivity within each module, known as hub genes. To pinpoint modules of interest that were significantly associated with the musical stimuli, module eigengenes were correlated with the time point data (bTP and fTP), and gene significance (GS), which measures the biological relevance of genes within the module, was assessed. For each gene, Module Membership (MM) indicates its intramodular connectivity within the module. Multiple test adjustments were performed using the FDR (False Discovery Rate) method according to the Benjamini-Hochberg procedure (Benjamini & Hochberg, 1995).

To analyze the functional aspects of the significantly correlated modules, an over-representation analysis was conducted using the *Clusterprofiler* R package (Wu et al., 2021). Additionally, biological processes from the Gene Ontology (GO) database were considered for this analysis. To make the results easier to interpret, redundant terms were identified and removed (similarity > 0.7) after computing the terms’ similarity matrix.

Different R packages were used to generate volcano plots (*EnhancedVolcano* (Blighe et al., 2020) and heatmaps (*ComplexHeatmap (Gu et al., 2016)*). Statistical significance was assessed using the Wilcoxon test. Statistical analyses were performed using R version (The R Core Team, 2019).

### Feature selection and predictive signature generation

The top 20 candidate biomarkers associated with musical stimulation in ASD were used to derive a predictive transcriptomic signature via a multi-algorithm feature-selection strategy. For this purpose, normalized gene expression data corrected for patient-specific effects from the discovery was used to train the models. The validation cohort was not used at any stage of feature selection or score construction and served exclusively for independent performance assessment.

To prevent information leakage arising from the paired samples, custom cross-validation folds were manually defined so that samples from the same patient (bTP and fTP) were always assigned to the same fold. To identify a robust biomarker signature, four machine learning algorithms available in the *caret* R package (Kuhn, 2008) machine learning algorithms were independently applied for feature selection using repeated cross-validation (5 folds repeated 30 times): *i*) Random Forest, *ii*) Elastic Net (*glmnet*), *iii*) Linear Support Vector Machine (SVM), and *vi*) Partial Least Squares (PLS). Model hyperparameters were optimized by maximizing the area under the receiver operating characteristic curve (AUC). The machine learning algorithms were exclusively used to identify robust biomarkers.

Because *glmnet*, SVM, and PLS are sensitive to differences in feature scale, predictors were centered and scaled within each cross-validation training fold using the preprocessing procedures implemented in caret. Model calibration was additionally assessed using the Brier score, calculated from the cross-validated predicted probabilities generated by the optimal hyperparameter configuration of each machine learning algorithm.

Variable importance was extracted from the final optimized model of each algorithm using the *varImp* function. For each algorithm, genes were ranked according to their variable importance scores. Individual rankings were integrated using the Robust Rank Aggregation (RRA) algorithm from the *RobustRankAggreg* R package (Kolde et al., 2012), producing a consensus ranking that identifies genes consistently prioritized across different machine learning approaches.

To reduce redundancy among highly correlated biomarkers, genes showing strong pairwise correlation (|*r*| > 0.80) were excluded, retaining the highest-ranked gene within each correlated group.

To obtain an interpretable diagnostic signature suitable for independent validation, candidate biomarkers were subsequently re-trained in the ASD discovery cohort using ridge-penalized logistic regression implemented in the *glmnet* R package (Friedman et al., 2010). The optimal regularization parameter (λ) was selected by internal cross-validation within the discovery cohort, and the resulting regression coefficients were used to define the per-sample score.

The coefficients obtained from the discovery cohort were applied directly to the independent validation datasets and HCs to calculate the score for each sample.

Diagnostic performance was assessed using receiver operating characteristic (ROC) analysis implemented with the *pROC* package (Robin et al., 2011). For each candidate signature, the following metrics were calculated: Area under the ROC curve (AUC), 95% confidence intervals, optimal threshold, sensitivity and specificity.

## Results

### Characteristics of the cohorts

The donors comprised individuals with ASD, HCs, and members of the general public. The Sensogenoma22 discovery cohort included 24 participants (median age: 18 years, range: 8-47; 37% female), comprising 11 individuals with ASD (median age: 16 years, range: 8-37; 36% female) and 13 HCs (median age: 19 years, range: 17-47; 38% female). The ASD cohort encompassed individuals with a broad spectrum of clinical presentations, including different levels of ASD severity and several neurodevelopmental or neurological comorbidities, such as attention-deficit/hyperactivity disorder (ADHD), intellectual disability, epilepsy, tuberous sclerosis, visual impairment, and rare genetic syndromes. Musical phenotypes were also heterogeneous: although only one participant had formal musical training, 10 of the 11 individuals self-reported at least one preserved musical ability, including good pitch perception, rhythmic abilities, accurate singing, or absolute pitch.

The Sensogenoma25 validation cohort included 14 participants, comprising 5 individuals with ASD (median age: 19 years, range: 12-28; 20% female) and 9 HCs (median age: 27 years, range: 18-48; 67% female). Similarly, the ASD validation cohort included participants with different clinical phenotypes, ranging from Asperger syndrome (ASD level 1) to ASD level 3, with associated comorbidities including ADHD, Williams syndrome, epilepsy, and acquired brain injury. Musical abilities were variable: two participants reported no musical skills, whereas three displayed preserved musical traits, including good pitch perception, rhythmic abilities, or formal musical training with absolute pitch.

Healthy controls from both cohorts represented a broad range of musical backgrounds, from individuals reporting no musical abilities to participants with elementary, professional, or non-formal musical training. Several controls played one or more musical instruments and reported musical traits such as absolute pitch, musical talent, creativity, accurate singing, good rhythmic abilities, or good pitch perception, providing substantial variability in musical aptitude across the control population.

### Transcriptomic response to music in the discovery cohort

The transcriptomic response to the musical intervention was first characterized in the 2022 discovery cohort by comparing saliva samples collected before (bTP) and after (fTP) the concert. In the ASD group, 520 genes were detected after filtering low-expressed genes, of which 33 genes were differentially expressed after multiple-testing correction (adjusted *P*-value < 0.05), including 23 upregulated and 10 downregulated genes (**Figure 2A**; **Table S1**). The largest expression changes (|log_2_FC| > 0.5) were observed for nine genes, comprising eight upregulated and one downregulated gene (**Table S1**). Among them, *PTGS2* and *CXCL8* exhibited the largest effect sizes (log_2_FC = 0.7).

**Figure 2.**
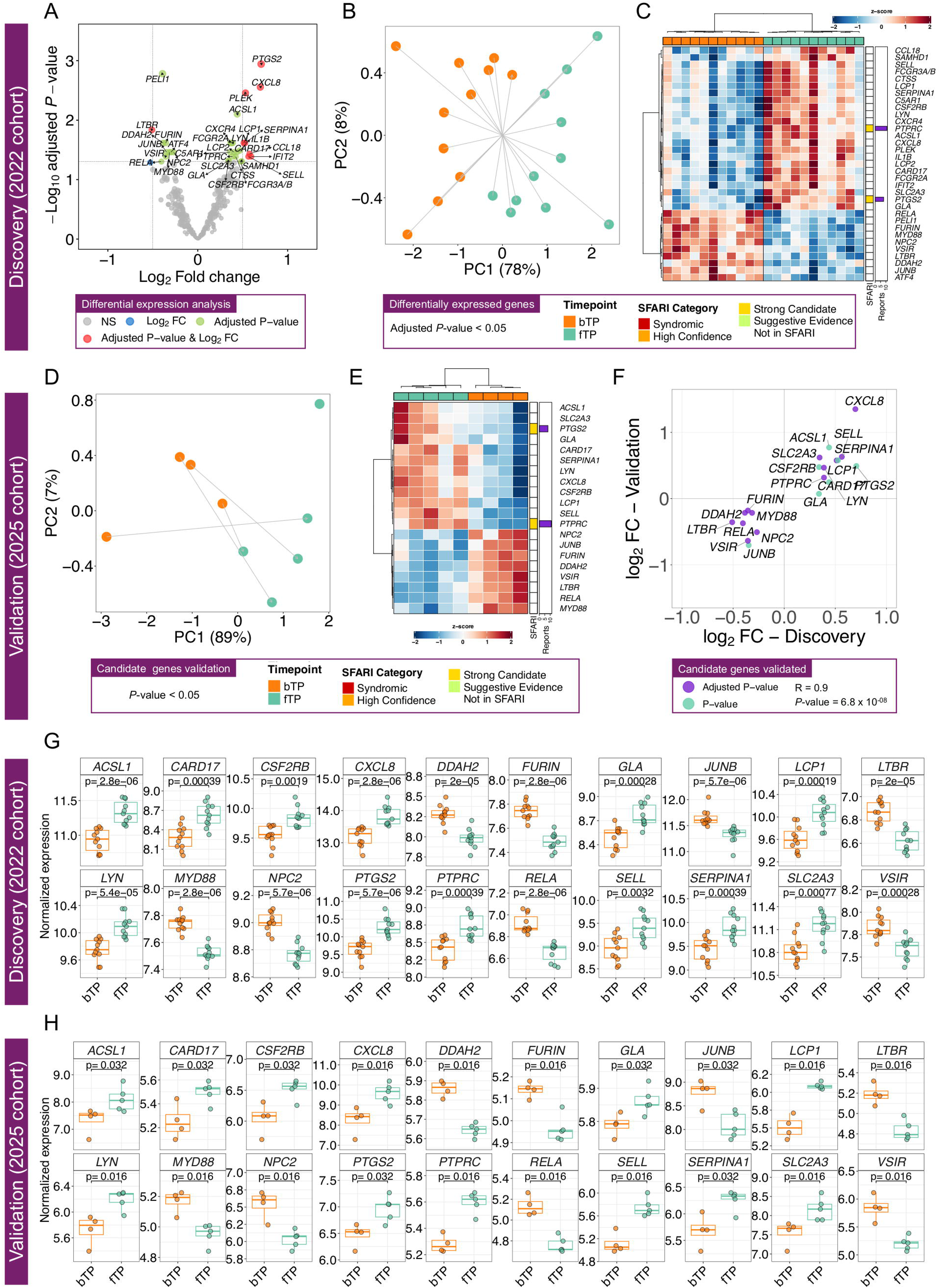
Transcriptomic response to the musical intervention in the discovery and validation ASD cohorts. (A) Volcano plot showing differential gene expression between baseline (bTP) and post-concert (fTP) samples in the 2022 discovery cohort. (B) Principal component analysis (PCA) based on the 33 DEGs identified in the discovery cohort. (C) Hierarchical clustering heatmap of the 33 DEGs. Genes are annotated according to their SFARI Gene evidence category. (D) PCA of the independent 2025 validation cohort based on the 20 validated candidate genes identified in the discovery cohort. (E) Hierarchical clustering heatmap of the validated candidate genes in the independent cohort. (F) Correlation of log_2_FC values between the discovery and validation cohorts, demonstrating a high concordance of transcriptional responses (Pearson’s *R* = 0.90, *P*-value = 6.8 × 10^-8^). (G) Boxplots showing normalized expression levels of the 20 DEGs in the discovery cohort subsequently validated in the validation cohort. (H) Boxplots showing normalized expression levels of the validated genes in the independent 2025 validation cohort.

To evaluate whether these DEGs detected in ASD patients collectively captured the transcriptional response to the musical intervention, PCA was performed using the expression levels of the 33 ASD DEGs. The first two principal components explained 78% and 8% of the total variance, respectively, and clearly separated bTP and fTP samples, demonstrating a coordinated transcriptional response across participants (**Figure 2B**).

The expression patterns of the same 33 DEGs showed a clear segregation of baseline and post-concert samples, with highly consistent expression profiles across individuals within each time point (**Figure 2C**). Cross-referencing the significant genes with the SFARI Gene database identified two ASD-associated genes within the music-responsive signature. Both *PTGS2* and *PTPRC* are currently classified as SFARI Gene score 2 (Strong Candidate genes), indicating prior evidence supporting their involvement in ASD.

In contrast to the ASD cohort, longitudinal analyses in HCs revealed only minimal transcriptional changes following musical stimulation, with only three genes (*CXCR4*, *HLA-DPA1*, and *HLA-DRA*) identified as differentially expressed (adjusted *P*-value < 0.05). All of the three genes were upregulated (log_2_FC >0.5) but showing marginal adjusted *P*-values (0.047; **Table S1**), indicating a substantially weaker transcriptomic response than that observed in ASD participants.

### Validation of candidate genes in an independent cohort

The 2025 cohort was subsequently used as an independent validation dataset to assess the reproducibility of the transcriptomic signature identified in the 2022 discovery cohort. Rather than performing a second transcriptome-wide differential expression analysis, only the 33 ASD DEGs identified in the 2022 discovery cohort were evaluated using a paired Wilcoxon signed-rank test.

Remarkably, 20 of the 33 candidate genes (60.6%) remained significantly differentially expressed (*P*-value < 0.05) in the independent validation cohort, despite its substantially smaller sample size and consequently lower statistical power (**Table S2**). Of these, 13 genes remained significant after multiple-testing correction (*CXCL8*, *DDAH2*, *FURIN*, *LCP1*, *LTBR*, *LYN*, *MYD88*, *NPC2*, *PTPRC*, *RELA*, *SELL*, *SLC2A3*, and *VSIR*), whereas an additional seven genes reached nominal significance (*ACSL1*, *CARD17*, *CSF2RB*, *GLA*, *JUNB*, *PTGS2*, and *SERPINA1*). Consistent with the findings in the discovery cohort, *CXCL8* also exhibited the largest effect size among the validated genes in the new independent cohort (log_2_FC = 1.35). Notably, of the two ASD-associated genes identified in the SFARI Gene database, *PTPRC* remained significant after multiple-testing correction in the validation cohort, whereas *PTGS2* was replicated at nominal significance.

At the global transcriptomic level, PCA based on the 20 validated genes revealed a clear separation between bTP and fTP samples, with the first two principal components explaining 89% and 7% of the total variance, respectively (**Figure 2D**). Similarly, hierarchical clustering of the same genes segregated baseline and post-concert samples, further supporting the reproducibility of the music-associated transcriptional signature in the independent cohort (**Figure 2E**).

To further quantify the reproducibility of the transcriptional response, the log_2_FC estimates obtained in the discovery and validation cohorts were compared for these 20 candidate genes. This analysis revealed a remarkably strong positive correlation (*R* = 0.90, *P*-value = 6.8 × 10^-8^), indicating that both the magnitude and direction of the transcriptional changes were highly consistent across independent cohorts with no discordant expression changes observed, further supporting the reproducibility of the identified transcriptomic signature after stimulation in ASD (**Figure 2F**). Boxplots of the 20 validated DEGs illustrate the individual expression values at bTP and fTP, together with the corresponding statistical significance, confirming the direction and magnitude of the transcriptional changes identified by the differential expression analysis in both the discovery (**Figure 2G**) and validation (**Figure 2H**) cohorts.

When analyzing the intermediate timepoint (iTP), no genes remained significantly differentially expressed between bTP and iTP after correction for multiple testing. Nevertheless, three genes (*FURIN*, *DDAH2*, and *PTPRC*) showed nominal significance (**Table S1**). Likewise, no statistically significant differences were detected between iTP and fTP, consistent with the heatmap clustering analysis, in which most iTP and fTP samples clustered together (**Table S1**, **Figure S1A**). Although no statistically significant differences were detected, visual inspection of the boxplots revealed apparent temporal expression patterns for several genes across the different time points, including, for instance, *CARD17*, *CSF3R*, *PTPRC*, and *FURIN* (**Figure S1B**). Interestingly, for most genes (80%) and despite limited sample size, median expression levels at the intermediate time point fell between those observed at the bTP and fTP (**Figure S1B**).

Similarly to the results observed in the HC’s discovery cohort, in the validation cohort none of the three candidate genes showed consistent differential expression. Although *HLA-DQA* reached nominal significance in both the discovery and validation cohorts between bTP and fTP with a concordant direction of effect, the magnitude of the change was modest (log_2_FC = 0.22) and did not remain significant after multiple-testing correction (**Table S2**).

The degree of transcriptomic replication observed in the independent validation cohort substantially exceeded that expected by chance. Under the specified null model, the number of replicated genes follows a binomial distribution, *X ∼ B*(33,0.05), with an expected value of *E*(*X*) = 1.65 genes. Instead, 20 of the 33 candidate genes remained significantly differentially expressed in the validation cohort, corresponding to a probability of only *P*(*X* ≥ 20) = 2.90 × 10^-18^. Furthermore, all replicated genes exhibited concordant directions of effect between cohorts. Considering both statistical significance and effect-direction concordance (*p* = 0.025), the expected number of replicated genes decreases to E(X) = 0.825, whereas the probability of observing 20 or more replicated genes is *Pr*(*X* ≥ 20) = 3.81 × 10^-24^. Assuming that the 33 genes identified in the discovery cohort represented a random subset of the 700 transcripts analyzed, the expected overlap under a purely random model would be approximately 33 × (33/700) = 1.56 genes, more than an order of magnitude lower than the 20 replicated genes observed in the validation cohort. The observed replication rate (20/33, 61%) was also markedly higher than the proportion of significant genes identified in the discovery phase (33/700, 4.7%), providing compelling statistical evidence that the observed replication is highly unlikely to be attributable to random variation and strongly supporting the robustness and reproducibility of the identified transcriptomic signature.

None of the replicated genes reached statistical significance in either the 2024 or 2025 HCs cohorts, despite control and ASD samples being processed simultaneously and subjected to identical experimental and analytical procedures.

### Enrichment analysis of the candidate genes altered in ASD

The Over-Representation Analysis (ORA) performed on the 20 validated genes revealed a highly coherent functional profile dominated by processes related to innate immunity, inflammation, and leukocyte activation (**Table S3**, **Figure 3**).

**Figure 3.**
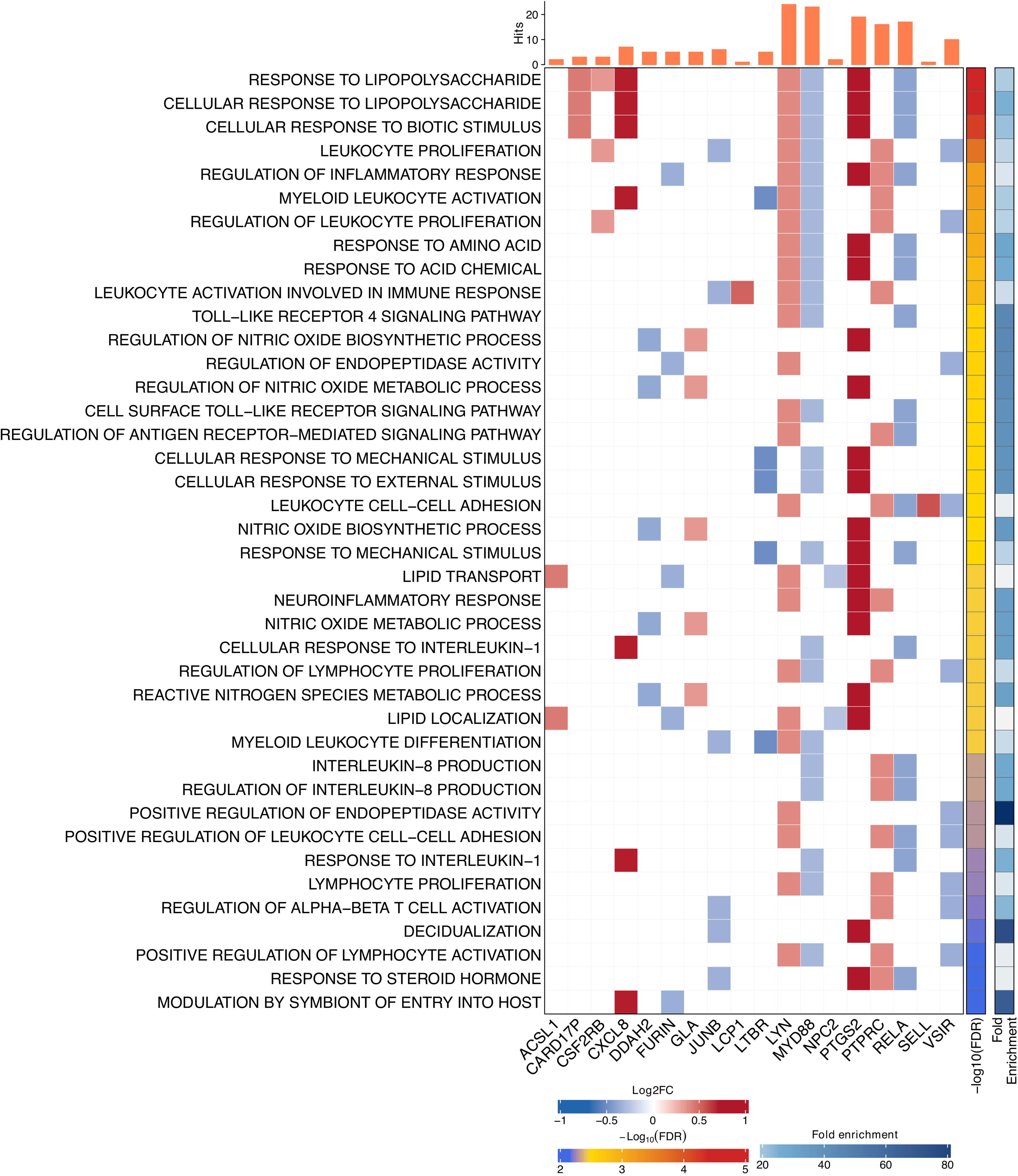
Top 40 significantly enriched Gene Ontology (GO) biological pathways identified by over-representation analysis of the 20 candidate genes modulated by musical stimulation and validated in the ASD cohort. Heatmap colors represent the log_2_FC of these genes obtained from the differential expression analysis (bTP *vs.* fTP) in the discovery cohort, adjusted *P*-values (shown as -log_10_ adjusted *P*-values), and fold enrichment of each pathway.

The most significantly enriched biological processes were associated with the response to lipopolysaccharide (LPS) and other microbial stimuli, including response to lipopolysaccharide, cellular response to lipopolysaccharide, and cellular response to biotic stimulus (**Table S3**, **Figure 3**). These enrichments were primarily driven by classical inflammatory genes such as *CXCL8*, *LYN*, *PTGS2*, and *CARD17*, indicating regulation of conserved host defense mechanisms.

Consistent with this inflammatory profile, multiple GO terms related to leukocyte activation, proliferation, and differentiation were significantly overrepresented, including leukocyte proliferation, regulation of leukocyte proliferation, leukocyte activation involved in immune response, myeloid leukocyte activation, myeloid leukocyte differentiation, lymphocyte proliferation, and positive regulation of lymphocyte activation. These processes involved several immune regulatory genes, including *PTPRC* (also known as *CD45*), *LYN*, and *MYD88*, suggesting coordinated modulation of both myeloid and lymphoid immune pathways (**Table S3**, **Figure 3**).

The analysis also identified significant enrichment of inflammatory signaling pathways, particularly the Toll-like receptor 4 (*TLR4*) signaling pathway, including Toll-like receptor 4 signaling pathway and cell surface Toll-like receptor signaling pathway, together with processes related to the response to interleukin-1 (*IL-1*), interleukin-8 production, and regulation of the inflammatory response. The prominent contribution of *MYD88*, *PTGS2*, and *CXCL8* highlights enrichment of canonical innate immune signaling pathways mediated through NF-κB (**Table S3**, **Figure 3**).

Another prominent functional category involved the metabolism and regulation of nitric oxide and reactive nitrogen species, with significant enrichment of terms such as nitric oxide biosynthetic process, regulation of nitric oxide metabolic process, and reactive nitrogen species metabolic process. These findings are consistent with modulation of antimicrobial and inflammatory effector mechanisms.

Additional enriched terms included cellular responses to mechanical and external stimuli, lipid transport and localization, leukocyte cell-cell adhesion, neuroinflammatory response, and modulation by symbiont of entry into host, suggesting that the transcriptional signature also encompasses processes involved in tissue remodeling, host-pathogen interactions, and environmental sensing (**Table S3**, **Figure 3**).

### Gene expression signature of musical stimulation in ASD

The 20 candidate biomarkers were independently evaluated for feature selection using four complementary machine learning algorithms. All machine learning approaches employed showed good calibration, with Random Forest and SVM exhibiting the best probability calibration (Brier scores of 0.027 and 0.022, respectively), whereas Elastic Net and PLS showed slightly higher prediction error (0.111 and 0.105, respectively).

The consensus ranking identified *FURIN*, *MYD88*, *RELA*, *JUNB* and *CXCL8* as the top five ranked biomarkers (**Table S4**, **Figure 4A**). *FURIN* was ranked first in the consensus ranking (score = 0.048) and was among the four most important variables in three out of the four machine learning algorithms used. To reduce the signature redundancy, only the highest-ranked gene within each correlated group was retained. This procedure yielded a final non-redundant panel of eight candidate genes (**Table S4**), with *FURIN*, *RELA*, *JUNB*, *CXCL8* and *LTBR* representing the top five highest-ranked biomarkers retained for signature construction. Signatures containing between two and five top-ranked genes added by ranking importance were evaluated for their potential to discriminate between bTP and fTP samples in the ASD discovery and independent validation cohorts. The minimal two-gene signature, composed of *FURIN* and *JUNB,* achieved perfect discrimination (AUC = 1.00; sensitivity = 100%; specificity = 100%) in both cohorts. Sequential addition of *RELA*, *CXCL8,* and *LBTR* did not improve the discriminatory performance of the two-gene signature (**Table S4**, **Figure 4B**).

**Figure 4.**
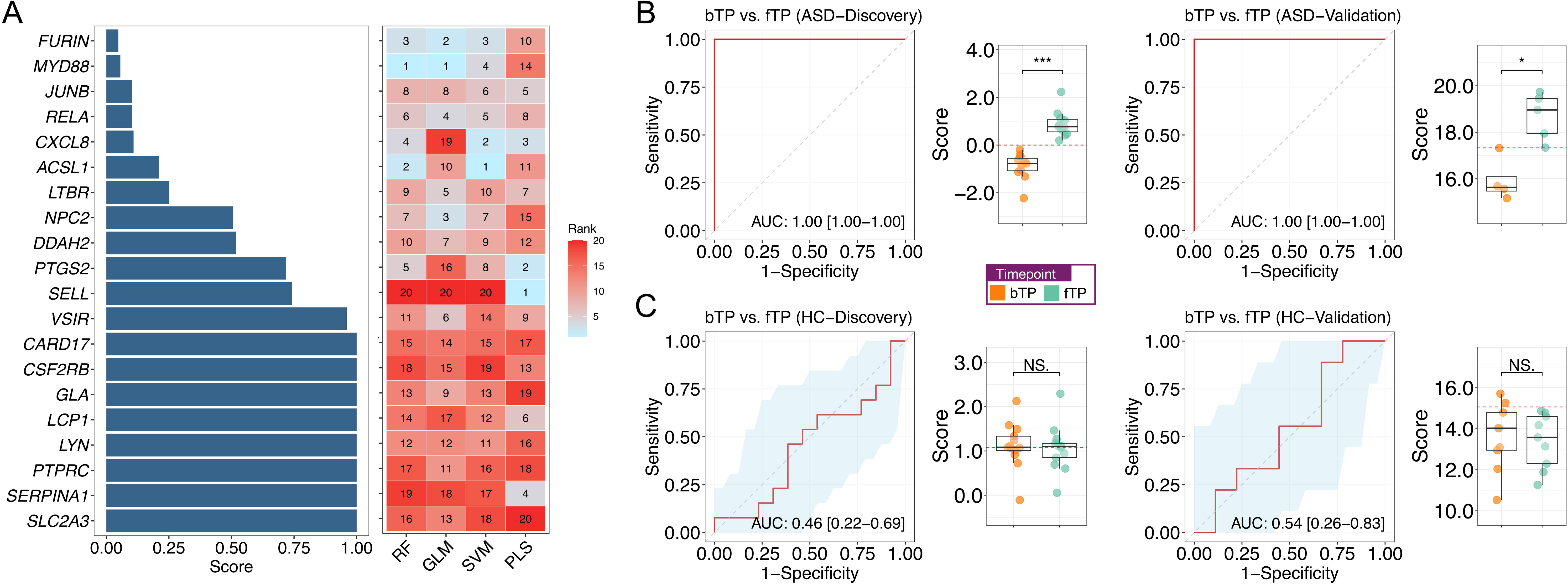
Gene-expression signature of musical stimulation in ASD. (A) Consensus ranking of candidate biomarkers identified by integration of four machine learning algorithms. Horizontal bars represent the Robust Rank Aggregation score for each candidate gene, with lower scores indicating greater agreement among algorithms. The heatmap shows the variable importance rank assigned by each machine learning model (RF: Random Forest, GLM: Elastic Net, SVM: Support Vector Machine, and PLS: Partial Least Squares). Numbers inside cells indicate the individual ranking of each gene. Diagnostic performance of the signature containing two genes (*FURIN*-*JUNB*) evaluated across the ASD discovery cohort and independent ASD validation cohort (B), and healthy control (HC) discovery and validation cohorts (C). Reported metrics include the area under the ROC curve (AUC) and 95% confidence interval (CI). Boxplots of the sample score values for each group are also represented. Two-sided Wilcoxon *P*-values are also represented by asterisks indicating the significance threshold (*** = 0.001, ** =0.01, * =0.05). The red line represents the optimal cut-off. Baseline time point: bTP, final time point: fTP.

To further assess their robustness, the signatures were evaluated using the iTP samples from the ASD validation cohort. The two- and three-gene signatures, comprising *FURIN*-*JUNB* and *FURIN*-*JUNB*-*RELA*, respectively, maintained perfect discrimination. In contrast, inclusion of additional genes (*CXCL8* and *LTBR*) resulted in a slight reduction in accuracy (AUC = 0.88, 95% CI: 0.59-1.00) (**Figure S2**, **Table S4**).

Performance differed substantially in the HCs cohorts; the discrimination power of the signatures in the HCs cohorts was very limited. In the discovery cohort, the three-gene signature (*FURIN*-*JUNB*-*RELA*) achieved the highest discrimination (AUC = 0.75, 95% CI: 0.53-0.96) (**Table S4**, **Figure 4C**). The remaining signatures showed AUC values ranging from 0.46 to 0.64. In the independent validation cohort, all signatures showed limited discrimination, with AUC values ranging between 0.42 and 0.54 (**Table S4**, **Figure 4C**).

### Co-expression modules correlated to musical stimulation

Weighted gene co-expression network analysis (WGCNA) identified distinct co-expression modules associated with the response to musical stimulation in the ASD and HC groups (**Table S5**).

In the ASD cohort, four co-expression modules were identified, two of the significantly associated with the bTP-fTP comparison after multiple-testing correction (**Figure 5A**; **Table S5**). The module represented by the hub gene *EIF3F* exhibited the strongest association, showing a significant negative correlation with the intervention (*r* = −0.836, *P*-value = 1.30 × 10^-6^, adjusted *P*-value = 6.51 × 10^-5^; **Figure 5A**; **Table S5**). In contrast, the module *FPR1* was positively associated with musical stimulation (*r* = 0.625, *P*-value = 1.89 × 10^-3^, adjusted *P*-value = 4.72 × 10^-3^).

**Figure 5.**
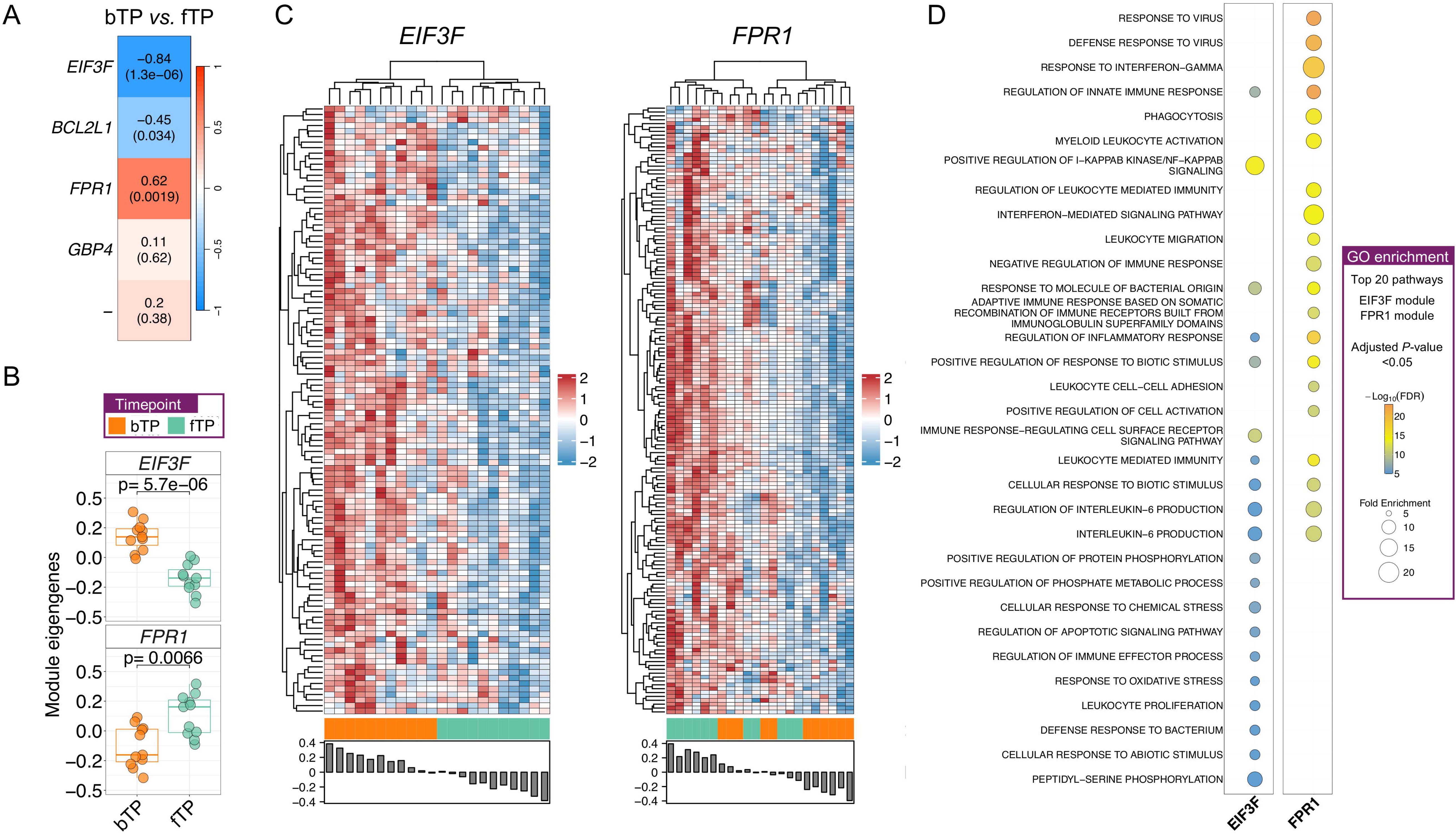
Co-expression network analysis in the ASD cohort. (A) Module-trait correlation heatmap showing the correlation between module eigengenes and the basal (bTP) versus post-stimulation (fTP) comparison. Correlation coefficients and corresponding *P*-values (in brackets) are indicated within each cell. (B) Distribution of per-sample module eigengene values for the EIF3F and FPR1 modules at bTP and fTP. Statistical significance was assessed using the Wilcoxon test. (C) Heatmaps showing the expression profiles of genes belonging to the EIF3F and FPR1 modules across all ASD samples. Bar plots below the heatmaps show the corresponding module eigengene values. (D) Functional enrichment analysis of the EIF3F and FPR1 modules. Bubble plots display the top 20 enriched Gene Ontology (GO) biological processes obtained for each module. Dot size represents the fold enrichment and associated with each GO term, and color indicates the adjusted P-value.

These associations were further supported by the distribution of module eigengenes across samples, with the *EIF3F* module exhibiting significantly lower eigengene values following musical stimulation (*P*-value = 5.7 × 10^-6^), whereas the *FPR1* module displayed the opposite trend, showing increased eigengene values after stimulation (*P*-value = 6.60 × 10^-3^); **Figure 5B**. Consistent with these findings, the expression profiles of genes belonging to the *EIF3F* module clearly differentiated basal and stimulated samples, reflecting a coordinated downregulation following musical stimulation, whereas genes within the *FPR1* module showed the opposite trend, with a more modest but coordinated upregulation after stimulation (**Figure 5C**). The *BCL2L1* module showed a moderate negative correlation associated with the musical stimulation (*r* = -0.45; *P-* value = 0.03), with the association remaining close to the threshold for statistical significance after multiple-testing correction (adjusted *P*-value = 0.056; **Table S5**).

Interestingly, the 20 candidate genes altered by musical stimulation in ASD represented core members of the *EIF3F* and *FPR1* co-expression modules, with many exhibiting high module membership (MM > 0.7) together with strong gene significance for the musical stimulation trait (**Figure S3**).

Functional enrichment analysis demonstrated that the two modules significantly associated with musical stimulation were characterized by distinct immune-related functional profiles, although they shared enrichment for several broad inflammatory processes. The negatively correlated *EIF3F* module was primarily enriched for biological processes related to NF-κB signaling and inflammatory regulation, whereas the positively correlated *FPR1* module was enriched for pathways involved in antiviral responses, interferon-γ signaling, and innate immunity (**Figure 5D**; **Table S6**). Notably, both modules were enriched for broader processes related to leukocyte-mediated immunity, IL-6-associated pathways, and inflammatory responses, indicating that these biological functions are represented by distinct but complementary co-expression networks (**Figure 5D**).

In the HC cohort, three different co-expression modules were detected, but only one of them reached statistical significance. The module represented by the hub gene *IFITM2* was positively correlated with the musical stimuli (*r* = 0.55, *P*-value = 3.25 × 10^-3^, adjusted *P*-value = 0.013 (**Figure S4A**; **Table S5**). Consistent with this significant correlation, both the module eigengene distribution and the expression profiles of genes within the *IFITM2* module showed a partial separation between bTP and fTP (**Figure S4B**-**C**). Over-representation analysis of the genes belonging to this module revealed that the top enriched biological processes were primarily related to antiviral responses, interferon-γ signaling, and T-cell proliferation and regulation (**Figure S4D**, **Table S6**).

## Discussion

A recent pilot study using saliva-based RNA sequencing in individuals with ASD provided the first evidence that music exposure can induce measurable molecular responses at multiple biological levels (Cavenaghi et al., 2025). By combining transcriptomic and exploratory metagenomic analyses, the study identified music-associated changes in genes and co-expression networks involved in neurodevelopment and immune regulation, as well as microbial alterations potentially relevant to neuroinflammation and ASD pathophysiology. Despite its limited sample size, these findings highlighted the potential of saliva-based transcriptomic profiling as a minimally invasive framework for monitoring biological responses to MIs in ASD. Building on these observations, the present study used a targeted transcriptomic approach in independent discovery and validation cohorts, enabling a more stringent assessment of candidate transcriptional response reproducibility.

One of the most remarkable findings was the marked contrast between the salivary transcriptional response to music in individuals with ASD and HCs. Whereas individuals with ASD exhibited a coherent and reproducible response across independent cohorts, HCs showed limited and diffuse transcriptional changes. This difference may reflect greater molecular sensitivity or responsiveness to the musical stimulus in ASD. Comparable findings were previously observed using a similar approach in neurodegenerative disorders (Gómez-Carballa et al., 2025). Moreover, the reproducibility of the ASD-associated response across independent cohorts suggests that this difference may not merely reflect a strong response in a subset of individuals, but rather a more consistently engaged biological response to musical stimulation in ASD.

Numerous behavioral and neuroimaging studies have reported superior perceptual processing, enhanced sensitivity to simple auditory features, and preserved or enhanced musical abilities in individuals with ASD (Chen et al., 2012; Chen et al., 2022; Gliga et al., 2015; Mottron et al., 2006; O’Riordan M, 2004; Ouimet et al., 2012). The present findings suggest that musical stimulation may engage sensory and associated neural circuits differently in ASD, potentially contributing to the more robust and consistent molecular response observed across two independent cohorts. Although this interpretation requires confirmation in larger cohorts, the reproducibility and cross-cohort concordance of the ASD-associated transcriptional signature are consistent with the hypothesis that musical stimulation elicits a distinct biological response in ASD compared with neurotypical individuals.

An important strength of the present study is the high reproducibility of the 33 genes initially identified in ASD individuals from the 2022 discovery cohort in an independent validation cohort from 2025. Despite the substantially smaller sample size of the validation cohort and reduced statistical power, 20 of these 33 candidate genes (61%) remained significantly differentially expressed, including 13 genes that withstood multiple-testing correction. The successful replication of these genes, together with the complete concordance in effect direction, supports the robustness and reproducibility of the transcriptional alterations despite the intrinsic heterogeneity in ASD. None of the replicated genes reached statistical significance in the 2024 or 2025 HCs cohorts, further reducing the likelihood of technical artifacts. These findings support the presence of an ASD-associated biological response to musical stimulation.

Notably, many of the replicated genes, including *CXCL8*, *MYD88*, *RELA*, *PTGS2*, *PTPRC*, *SELL*, *LYN*, and *LTBR*, are central regulators of innate immune signaling and inflammatory responses. Among the validated genes in ASD, *CXCL8* (*IL-8*) exhibited the largest effect size in both the discovery and validation cohorts. *CXCL8* is a pleiotropic chemokine involved in neutrophil recruitment, innate immune activation, and neuroimmune communication, and has emerged as one of the inflammatory mediators most consistently associated with ASD. Several studies have reported elevated circulating IL-8 levels in plasma from ASD individuals, with higher concentrations correlating with higher impairments in communication, social functioning, and behavioral symptoms (Ashwood et al., 2011; Shen et al., 2021). Although not all studies have replicated these associations (Anastasescu et al., 2024; Han et al., 2017), recent evidence continues to support a role for *CXCL8* in the immune dysregulation underlying ASD and highlights its potential utility as a biomarker of neuropsychiatric disorders and disease-related inflammatory processes (Masi et al., 2017; Shkundin & Halaris, 2024). The robust and reproducible upregulation of *CXCL8* observed following musical stimulation therefore suggests that this chemokine may participate in modulating the neuroimmune response elicited by the intervention in ASD.

The identification of *PTGS2* and *PTPRC* among the replicated music-responsive genes is noteworthy given the growing evidence implicating both genes in ASD. *PTGS2*, which encodes cyclooxygenase-2 (COX-2), is a key mediator of prostaglandin synthesis and inflammatory signaling. A genetic association between *PTGS2* and ASD has previously been reported in a Korean population cohort (Yoo et al., 2008), and *PTGS2* knockout mice display several autism-related phenotypes, including increased repetitive behaviors and impaired social interactions (Wong et al., 2019). Similarly, *PTPRC* encodes CD45, an essential regulator of T- and B-cell receptor signaling and immune homeostasis. Rare deletions involving *PTPRC* have been identified in individuals with ASD (Pinto et al., 2010), and a missense variant segregating within an extended multiplex ASD family was subsequently reported (Cukier et al., 2014). The presence of these two ASD-associated genes within the replicated music-responsive transcriptional signature is particularly intriguing because both converge on immune and inflammatory pathways implicated in ASD pathophysiology. Although the present findings do not establish a causal relationship between music exposure and ASD-related molecular mechanisms in the buccal cavity, they are consistent with music-induced transcriptional responses involving biological pathways that overlap with processes previously associated with the disorder, thereby supporting the biological plausibility of the identified response.

The enrichment analysis demonstrates that the 20 candidate genes involved in the transcriptomic response to music in ASD individuals converge on a robust pro-inflammatory and innate immune signature, characterized by activation of TLR/IL-1-MYD88 signaling, leukocyte activation and proliferation, production of inflammatory mediators, and nitric oxide-dependent antimicrobial responses. The predominance of innate immune and inflammatory pathways is consistent with accumulating evidence implicating immune dysregulation and neuroinflammation in ASD (Estes & McAllister, 2015; Gandal et al., 2022; Hughes et al., 2023). Their coordinated modulation following musical stimulation suggests that music may influence biological processes implicated in ASD through regulation of neuroimmune pathways.

The buccal transcriptomic response to musical stimulation in individuals with ASD can be accurately captured by a minimal gene signature derived from the panel of 20 validated candidate genes. Although all signatures comprising two to five genes exhibited excellent discriminatory performance, the minimal two-gene model, consisting of *FURIN* and *JUNB*, consistently matched the performance of the more complex models in both the discovery and independent validation ASD cohorts. Notably, the same signature accurately discriminated baseline and intermediate samples collected after only 25 minutes of music exposure in the validation cohort (detected in the iTP), indicating that the molecular response is already detectable at an early stage of the intervention. Moreover, the expression levels of most signature genes at the intermediate time point were intermediate between baseline and final values, consistent with a progressive, time-dependent transcriptional response to musical stimulation. In contrast, these signatures showed poor discriminatory ability in HCs, suggesting that they capture a molecular response specific to individuals with ASD. From a translational perspective, these findings support the potential use of these signatures as objective molecular biomarkers for monitoring responses to MIs. Such biomarkers could provide a molecular framework for comparing different therapeutic protocols, identifying stronger biological responders, and defining molecular endpoints for future clinical trials. Moreover, the small size of the signature and its development using buccal samples further enhance its translational potential, particularly in ASD individuals. Buccal sampling is minimally invasive and readily implemented using widely available targeted gene expression platforms, such as RT-qPCR, digital PCR, or NanoString.

Co-expression analysis further indicates that this response involves coordinated remodeling of transcriptional networks rather than isolated changes in gene expression in individuals with ASD. Two independent modules exhibited opposite responses to musical stimulation: the *EIF3F* module showed coordinated downregulation of genes involved in NF-κB-dependent inflammatory signaling, whereas the *FPR1* module showed increased expression of genes related to antiviral responses, interferon-γ signaling, and innate immunity. Although both modules were enriched for common broad immune-related processes, they represented distinct transcriptional programs through which musical stimulation could potentially reshape complementary aspects of the immune response. This interconnected mechanism is consistent with models proposing that immune dysregulation in ASD reflects complex alterations across multiple interconnected immune pathways (Estes & McAllister, 2015; Hughes et al., 2018). Interestingly, the hub gene *EIF3F* encodes a subunit of the mammalian eIF3 (eukaryotic translation initiation factor 3 subunit F) and has been reported in SFARI as a gene with suggestive evidence of association with ASD. Recent genetic studies have implicated *EIF3F* in ASD and other developmental disorders through pathogenic variants, providing genetic evidence of its potential role in ASD susceptibility (Huffmeier et al., 2021; Lakatosova et al., 2024; Lob et al., 2025; Martin et al., 2018; Repiska et al., 2025). It The validated candidate genes occupied central positions within the *EIF3F* and *FPR1* modules, indicating that the genes identified are not isolated transcriptional changes but key components of coordinated regulatory programs.

In contrast, the transcriptomic response in HCs was characterized by a single significant co-expression module centered on the *IFITM2* gene, predominantly enriched for antiviral and interferon-related pathways as well as T-cell-related processes. These processes partially overlap with those identified in the positively correlated FPR1 module in ASD, pointing to a common response of immune programs potentially representing conserved transcriptional responses to musical stimulation.

Several limitations should be considered when interpreting the present findings. First, although the sample size remains relatively modest, this limitation should be interpreted in the context of the substantial challenges involved in performing molecular studies in ASD. The recruitment of well-characterized participants, the implementation of standardized music exposure protocols, and the collection of high-quality biological samples under controlled conditions are particularly demanding in neurodevelopmental disorders characterized by marked clinical heterogeneity. The inclusion of an independent validation cohort represents a major strength, and, despite its limited sample size, it substantially increases the robustness of the identified molecular signature and reduces the likelihood that the findings reflect cohort-specific effects. Second, saliva presents unique challenges for transcriptomic analyses because the abundance of microbial RNA can interfere with host RNA sequencing, making unbiased whole-transcriptome approaches technically demanding. To overcome this limitation, a targeted NanoString gene expression panel was employed, enabling direct digital quantification of selected host transcripts without amplification and avoiding interference from the oral microbiota.

Finally, despite the technical challenges, saliva represents a particularly attractive biological matrix for studies involving individuals with ASD. Its non-invasive and well-tolerated collection facilitates participant recruitment and repeated sampling, making it especially suitable for pediatric and neurodevelopmental populations, where blood collection may be difficult or distressing. Importantly, a previous saliva-based RNA-sequencing study (Cavenaghi et al., 2025) demonstrated the feasibility of simultaneously characterizing host transcriptomic and microbial responses to music exposure in ASD, highlighting the potential of saliva as a minimally invasive biospecimen for longitudinal molecular monitoring and precision intervention studies in neurodevelopmental disorders.

The findings support the use of saliva-based molecular profiling as a scalable platform for investigating individual variability in responses to music and other sensory interventions. Future studies integrating molecular, behavioral, physiological, and clinical measures may help determine whether these responses can be used to identify biologically distinct subgroups and guide more personalized intervention in ASD.

## Ethics approval and Consent to participate

Written informed consent was obtained from the legal guardians or responsible parties of all participating individuals. The study was approved by the Ethics Committee of the Xunta de Galicia (registration code: 2020/021) and conducted in accordance with the principles outlined in the Declaration of Helsinki.

## Consent for publication

All participants have given permission to the publication of the project’s findings.

## Data availability statement

The raw count data from the human-merged and the metagenomic-merged datasets, along with the metadata, are publicly available in the Figshare repository under the accession: ##### (**Note**: This link will remain unavailable until the manuscript is published. In the meantime, and for review purposes, the data can be accessed using the following private token: ####).

## Author’s contribution

ASE, LN, and FMT conceived the project and funding. AC, AGC, ASE, and NeZM carried out the analyses. AGC, ASE, and NeZM contributed to a critical discussion of the main findings and wrote the first draft of the study. All the authors critically review the final version of the manuscript.

## Acknowledgements

The authors would like to express their appreciation to the study investigators of the Sensogenomics network (sensogenomics.com; Sensogenomics Working Group [see Annex]), as well as the nursery and laboratory service at the Hospital Clínico Universitario de Santiago de Compostela, for their invaluable dedication and support. This research project was made possible through the access granted by the Galician Supercomputing Center (CESGA) to its supercomputing infrastructure. The supercomputer FinisTerrae III and its permanent data storage system have been funded by the Spanish Ministry of Science and Innovation, the Galician Government, and the European Regional Development Fund (ERDF). We extend our deepest gratitude to the Sensogenomics Working Group (see annex for details) for their meaningful involvement and support throughout this project. The authors declare that they have no conflict of interest.

## Legend to the supplementary figures and tables

Table S1. DEGs after musical stimulation in ASD individuals and healthy controls. **Table S2.** Validation of candidate genes in the validation cohort. Validation of 33 DEGs identified in the ASD and discovery cohort (2022) and 14 DEGs (nominal *P*-value< 0.05) identified in the healthy controls (HC) discovery cohort (2022) using an independent validation cohort (2025). *P*-values were calculated using the Wilcoxon test. Genes showing an adjusted *P*-value <0.05 were highlighted in blue. bTP: basal timepoint; iTP: intermediate timepoint; fTP: final timepoint.

Table S3. Over-representation analysis from the 20 DEGs validated in the 2025 cohort. GO: Gene Ontology.

Table S4. Ranking of the twenty candidate genes according to their variable importance across four machine learning algorithms: Random Forest (RF), Elastic Net (GLM), Support Vector Machine (SVM), and Partial Least Squares (PLS) (upper panel). The last column indicates whether each gene was retained after correlation filtering. Diagnostic performance of the different gene signatures containing two to five genes evaluated across the ASD discovery cohort, independent ASD validation cohort, and healthy control (HC) discovery and validation cohorts (lower panel). Reported metrics include the area under the ROC curve (AUC), 95% confidence interval (CI), optimal classification threshold, specificity, and sensitivity. Baseline time point: bTP, final time point: fTP, intermediate time point: iTP.

Table S5. Co-expression module analysis in ASD patients and healthy controls (HC).

Table S6. Over-representation analysis for GO terms from statistically significantly correlated module with musical stimuli in ASD individuals and healthy controls (HC). GO: Gene Ontology

Figure S1. (A) Hierarchical clustering heatmap of the validated candidate genes in the independent cohort. (B) Boxplots showing normalized expression levels of the validated genes in the independent 2025 cohort. Statistical significance was assessed using the Wilcoxon test. bTP: basal timepoint; iTP: intermediate timepoint; fTP: final timepoint.

Figure S2. Diagnostic performance of the 2-RNA transcript signature containing (*FURIN* and *JUNB* genes) evaluated in the ASD validation cohort comparing the basal time point (bTP) with the intermediate time point (iTP). Reported metrics include the area under the ROC curve (AUC) and 95% confidence interval (CI). Boxplots of the sample score values for each group are also represented. Two-sided Wilcoxon *P*-values are also represented by asterisks indicating the significance threshold (*** = 0.001, ** = 0.01, * = 0.05). The red line represents the optimal cut-off.

Figure S3. Scatter plots showing the correlation between absolute module membership (MM) and absolute gene significance (GS) for all genes within the *EIF3F* (A) and *FPR1* (B) modules. Candidate genes validated across the discovery and validation cohorts are highlighted and labeled in italics. Dashed lines indicate the thresholds (MM > 0.7; GS > 0.5). Spearman’s correlation coefficient (ρ) and corresponding *P*-value are shown for each module.

Figure S4. Co-expression network analysis in the HC cohort. (A) Module-trait correlation heatmap showing the correlation between module eigengenes and the basal (bTP) versus post-stimulation (fTP) comparison. Correlation coefficients and corresponding *P*-values (in brackets) are indicated within each cell. (B) Distribution of per-sample module eigengene values for the IFITM2 module at bTP and fTP. Statistical significance was assessed using the Wilcoxon test. (C) Heatmaps showing the expression profiles of genes belonging to the IFITM2 module across all ASD samples. Bar plots below the heatmaps show the corresponding module eigengene values. (D) Functional enrichment analysis of the IFITM2 module. Bubble plot displays the top 20 enriched Gene Ontology (GO) biological processes obtained. Dot size represents the fold enrichment and is associated with each GO term, and color indicates the adjusted *P*-value.

## Sensogenomics Working Group

Antonio Salas Ellacuriaga - PI; Federico Martinón-Torres - PI; Laura Navarro Ramón - Coordinator *GenPoB/GenVip - Instituto de Investigación Sanitaria (IDIS) (alphabetic order)*

Alba Camino Mera, Albert Padín Villar, Alberto Gómez Carballa, Alejandro Pérez López, Alicia Carballal Fernández, Ana Cotovad Bellas, Ana Isabel Dacosta Urbieta, Ana María Pastoriza Mourelle, Ana María Senín Ferreiro, Andrés Muy Pérez, Antía Rivas Oural, Antonio Justicia Grande, Antonio Piñeiro García, Anxela Cristina Delgado García, Belén Mosquera Pérez, Blanca Díaz Esteban, Carlos Durán Suárez, Carmen Curros Novo, Carmen Gómez Vieites, Carmen Rodríguez-Tenreiro Sánchez, Celia Varela Pájaro, Claudia Navarro Gonzalo, Cristina Serén Trasorras, Cristina Talavero González, Einés Monteagudo Vilavedra, Estefanía Rey Campos, Esther Montero Campos, Fernando Álvez González, Fernando Caamaño Viñas, Francisco García Iglesias, Gloria Viz Rodríguez, Hugo Alberto Tovar Velasco, Irene Álvarez Rodríguez, Irene García Zuazola, Irene Rivero Calle, Iria Afonso Carrasco, Isabel Ferreirós Vidal, Isabel Lista García, Isabel Rego Lijo, Iván Prieto Gómez, Iván Quintana Cepedal, Jacobo Pardo Seco, Jesús Eirís Puñal, José Gómez Rial, José Manuel Fernández García, José María Martinón Martínez, Julia Cela Mosquera, Julia García Currás, Julián Montoto Louzao, Lara Martínez Martínez, Laura Navarro Marrón, Lidia Piñeiro Rodríguez, Lorenzo Redondo Collazo, Lúa Castelo Martínez, Lucía Company Arciniegas, Luis Crego Rodríguez, Luisa García Vicente, Manuel Vázquez Donsión, María Dolores Martínez García, María Elena Gamborino Caramés, María Elena Sobrino Fernández, María José Currás Tuala, María Martínez Leis, María Soledad Vilas Iglesias, María Sol Rodriguez Calvo, María Teresa Autran García, Marina Casas Pérez, Marta Aldonza Torres, Marta Bouzón Alejandro, Marta Lendoiro Fuentes, Miriam Ben García, Miriam Cebey López, Montserrat López Franco, Narmeen Mallah, Natalia García Sánchez, Natalia Vieito Perez, Nour El Zahraa Mallah, Patricia Regueiro Casuso, Ricardo Suárez Camacho, Rita García Fernández, Rita Varela Estévez, Rosaura Picáns Leis, Ruth Barral Arca, Sandra Carnota Antonio, Sandra Viz Lasheras, Sara Pischedda, Sara Rey Vázquez, Sonia Marcos Alonso, Sonia Serén Fernández, Susana Rey García, Vanesa Álvarez Iglesias, Victoria Redondo Cervantes, Vanesa Álvarez Iglesias, Wiktor Dominik Nowak, Xabier Bello Paderne, Xabier Mazaira López

### Nursing volunteers (alphabetic order)

Alejandra Fernández Méndez, Ana Isabel Abadín Campaña, Ana María León Caamaño, Ana María Buide Illobre, Ángeles Mera Cores, Carmen Nieves Vastro, Carolina Suarez Crego, Concepción Rey Iglesias, Cristina Candal Regueira, Dolores Barreiro Puente, Elvira Rodríguez Rodríguez, Eugenia González Budiño, Eva Rey Álvarez, Fernando Rodríguez Gerpe, Gemma Albela Silva, Isabel Castro Pérez, Isabel Domínguez Ríos, José Ángel Fernández de la Iglesia, José Cruces Vázquez, José Luis Cambeiro Quintela, José Ramón Magariños Iglesias, Julia Rey Brandariz, Julio Abel Fernández López, Luisa García Vicente, Manuel González Lito, Manuel González Lijó, Manuela Pérez Rivas, Margarita Turnes Paredes, María Aurora Méndez López, María Begoña Tomé Arufe, María Campos Torres, María del Carmen Baloira Nogueira, María del Carmen García juan, María Esther Moricosa García, María Luz Chao Jarel, María Martínez Leis, María Mercedes Jiménez Santos, María Salomé Buide Illobre, María Victoria López Pereira, Mercedes Jorge González, Mercedes Isolina Rodríguez Rodríguez, Miren Payo Puente, Natalia Carter Domínguez, Olga María Reyes González, Pilar Mera Rodríguez, Purificación Sebio Brandariz, Salomé Quintáns lago, Yolanda Rodríguez Taboada, María Pereira Grau.

### Other volunteers (alphabetic order)

Alba Arias Gómez, Alejandro Moreno Díaz, Ana Arca Marán, Astro González Guirado, Brais García Iglesias, Carlos Sánchez Rubín, Carmen Otero de Andrés, Clara Pérez Errazquin Barrera, Claudia Rey Posse, Cristina Rojas García, Eduardo Xavier Giménez Bargiela, Elena Gloria Morales García, Fabio Izquierdo García Escribano, Gabriel Guisande García, Jaime López Martín, Lara Pais Ramiro, Lucía Rico Montero, Luís Estévez Martínez, Manuel Estévez Casal, María Aránzazu Palomino Caño, María Rubio Valdés, Marisol Nogales Benítez, Miryam Tilve Pérez, Nuria Villar Muiños, Pablo Del Cerro Rodríguez, Pablo Pozuelo Martínez Cardeñoso, Salma Ouahabi El Ouahabi, Santiago Vázquez Calvach

